# The scale-invariant, temporal profile of neuronal avalanches in relation to cortical γ–oscillations

**DOI:** 10.1101/757278

**Authors:** Stephanie R. Miller, Shan Yu, Dietmar Plenz

**Affiliations:** Section on Critical Brain Dynamics, National Institute of Mental Health, Bethesda, MD, USA; Inst. for Physical Science and Technology, Univ. of Maryland College Park, College Park, MD, USA; Brainnetome Center, Inst. of Automation, Chinese Academy of Sciences, China

**Author notes:** Correspondence: Dietmar Plenz, Ph.D., Section on Critical Brain Dynamics, National Institute of Mental Health, Porter Neuroscience Research Center, Rm 3A-1000, 35 Convent Drive, Bethesda, MD 20892.

**Keywords:** Nonhuman primate, cortex, local field potential, resting activity, neuronal avalanches, oscillations, criticality, high-density microelectrode arrays

## Abstract

Activity cascades are found in many complex systems. In the cortex, they arise in the form of neuronal avalanches that capture ongoing and evoked neuronal activities at many spatial and temporal scales. The scale-invariant nature of avalanches suggests that the brain is in a critical state, yet predictions from critical theory on the temporal unfolding of avalanches have yet to be confirmed in vivo. Here we show in awake nonhuman primates that the temporal profile of avalanches follows a symmetrical, inverted parabola spanning up to hundreds of milliseconds. This parabola constrains how avalanches initiate locally, extend spatially and shrink as they evolve in time. Importantly, parabolas of different durations can be collapsed with a scaling exponent close to 2 supporting critical generational models of neuronal avalanches. Spontaneously emerging, transient γ–oscillations coexist with and modulate these avalanche parabolas thereby providing a temporal segmentation to inherently scale-invariant, critical dynamics. Our results identify avalanches and oscillations as dual principles in the temporal organization of brain activity.

**Significance Statement:** The most common framework for understanding the temporal organization of brain activity is that of oscillations, or ‘brain waves’. In oscillations, distinct physiological frequencies emerge at well-defined temporal scales, dividing brain activity into time segments underlying cortex function. Here, we identify a fundamentally different temporal parsing of activity in cortex. In awake Macaque monkeys, we demonstrate the motif of an inverted parabola that governs the temporal unfolding of brain activity in the form of neuronal avalanches. This symmetrical motif is scale-invariant, that is, it is not tied to time segments, and exhibits a scaling exponent close to 2, in line with prediction from theory of critical systems. We suggest that oscillations provide a transient regularity in an otherwise scale-invariant temporal organization pervading cortical activity at numerous scales.

## INTRODUCTION

Cascades are found in many complex systems. The dissemination of information in social media^1^, the spread of infections during epidemics^2^, and the propagation of neuronal activity in the brain^3^ are prominent examples of how cascades and cascade failures^4–6^ provide insight into system function. In the brain, cascading activity has been identified in the form of neuronal avalanches by the presence of power laws in the distributions of avalanche size and duration^3, 7, 8^. The scale-invariant nature of neuronal avalanches describes the firing of nerve cells^9, 10^ and, at larger scales, captures neuronal population dynamics in zebrafish^11^, nonhuman primates^7, 12, 13^, and humans^14–19^. This supports the idea that the brain might operate close to a critical state^8, 20–22^ where numerous aspects of information processing are optimized^23–30^. Yet, avalanche size and duration statistics do not provide insight into the process of cascading itself. This process can be more rigorously assessed by studying the temporal profile of avalanches, that is, how avalanches initiate locally and expand and shrink spatially as they evolve in time.

Theory and simulations^31–34^ predict that cascade profiles follow an inverted parabola in critical systems, but it is currently not known whether such a unique avalanche profile guides brain activity. Variable and asymmetric profiles have been reported for neuronal cultures^35, 36^, and profiles seem to depend on avalanche duration in humans^18^. Identifying the correct profile of neuronal avalanches will provide insights into the temporal evolution of brain activity, will distinguish between different models of avalanche generation^37–40^ and, importantly, might provide a biomarker given recent findings that profiles predict recovery from brain insults^41, 42^. A second prediction from critical theory is that the avalanche parabola can be collapsed over many avalanche durations with a scaling exponent, χ, larger than 1.5 (Refs.^31–34^). Robust collapses, though, require a large amount of data to include long avalanches, which are rare. Equally challenging are recent reports that χ ranges between 1–1. 5 for rodent tissue *in vitro*^35, 36^ and *in vivo*^43^, as well as in human MEG^18^, suggestive of non-critical dynamics^38^. A scaling collapse with χ ≅ 2 was recently found in zebrafish whole brain activity^11^, yet it is unknown whether this is the case for mammalian cortex.

Compared to non-biological systems, profile estimates in the brain are further challenged by the presence of prominent brain oscillations, such as gamma activity (γ; ∼30–100 Hz)^44–46^. Oscillations establish temporal scales for neuronal dynamics with activity arising around specific cycle times^47–49^. Such a scale-dependent temporal organization based on oscillations seems contradictory to the scale-invariant temporal organization encountered in avalanches. Specifically, it is currently not known how the scale introduced by an oscillation affects the temporal evolution of avalanches.

Here we reliably identify avalanche profiles at several temporal resolutions, 1–30 ms, using long-term recordings from awake nonhuman primates with high-density microelectrode arrays. By overcoming statistical and oscillation-induced constraints in profile collapse, we show that neuronal avalanches exhibit the scale-invariant profile of an inverted parabola with a scaling exponent χ ≅ 2, as predicted for critical dynamics. We also demonstrate that the removal of γ– oscillations by bandpass filtering abolishes the critical avalanche profile thus demonstrating interdependence and coexistence of critical dynamics during intermittent oscillation periods. Our findings suggest a novel scale-invariant temporal motif that governs neuronal activity in cortex over many durations to which γ-oscillations add distinct regularity in time.

## RESULTS

### Neuronal avalanches and oscillations coexist in ongoing brain activity

We studied the temporal profile of neuronal avalanches and its potential interdependence on oscillations in the ongoing local field potential (LFP) of nonhuman primates. Using chronically implanted high-density microelectrode arrays, the ongoing LFP (1–100 Hz) was recorded in premotor (PM, *n* = 2) and prefrontal cortex (PF, *n* = 4) of three nonhuman primates (*Macaca mulatta*; K, V, N) sitting in a monkey chair. The animals were awake during the recording sessions but were not engaged in behavioral tasks. For each array, about 4 **±** 2 hr (mean ± s.d.) of activity were analyzed during 9 ± 7 recording sessions over the course of 5 ± 4 weeks (85 ± 8 electrodes/array; Supplementary Table 1; see Methods).

Avalanches were identified first by extracting from each electrode negative LFP transients (nLFPs) exceeding a threshold of −2 s.d. (Fig. 1a). nLFPs occurred at a rate of 3.4 ± 0.3 Hz per electrode and were separated on the array by an average inter-event interval, ⟨IEI⟩ = 3.4 ± 0.6 ms (Supplementary Table 2; *n* = 6 arrays). In line with experiments^3, 12^ and theory^50^, we then concatenated successive time bins with nLFPs on the array at temporal resolution Δt = ⟨IEI⟩ into nLFP clusters (Fig. 1b). We found that nLFP cluster size, S, i.e. the number of nLFPs in each cluster, distributed as a power law P(S) ∼ S^−α^ with a slope α ≅ 3/2, the hallmark of neuronal avalanches (Fig. 1c; power law vs. exponential, LLR>>10^4^, *P* < 0.001; Ref.^51, 52^). In contrast, avalanche durations, T, did not distribute according to a power law P(T) ∼ T^−β^ and no duration slope β could be obtained at Δt = ⟨IEI⟩ (Fig. 1d; LLR < 0; *P* > 0.1). Instead, durations T, or, alternatively, lifetimes L = T/Δt, distributed shorter than expected and were closer to an exponential distribution. This suggested the presence of a secondary process affecting avalanche durations. Indeed, the average periodogram of the continuous LFP, besides having a 1/f*^a^* decay with *a* ≅ 1 (Ref.^53^), indicated a prominent oscillation at 29.3 ± 1.7 Hz (*n* = 6 arrays; Fig. 1e). Oscillations between 25–35 Hz are within the lower regime of the γ–band^46^, but they have also been identified as the β2 sub-band in primate PF and PM^45, 54^. These oscillations, which we will refer to simply as γ–activity, were variable in peak frequency and power, spatially heterogeneous and differed significantly between arrays and areas (Fig. 1f; *P* < 0.001, ANOVA, *F_power_* = 133.6, *F_frequency_* = 86.1). Establishing the coexistence of avalanches and γ–oscillations in ongoing activity of motor and prefrontal cortex in nonhuman primates paved the way to study the interdependencies regarding critical measures of avalanche dynamics and oscillation strength.

**Fig. 1:**
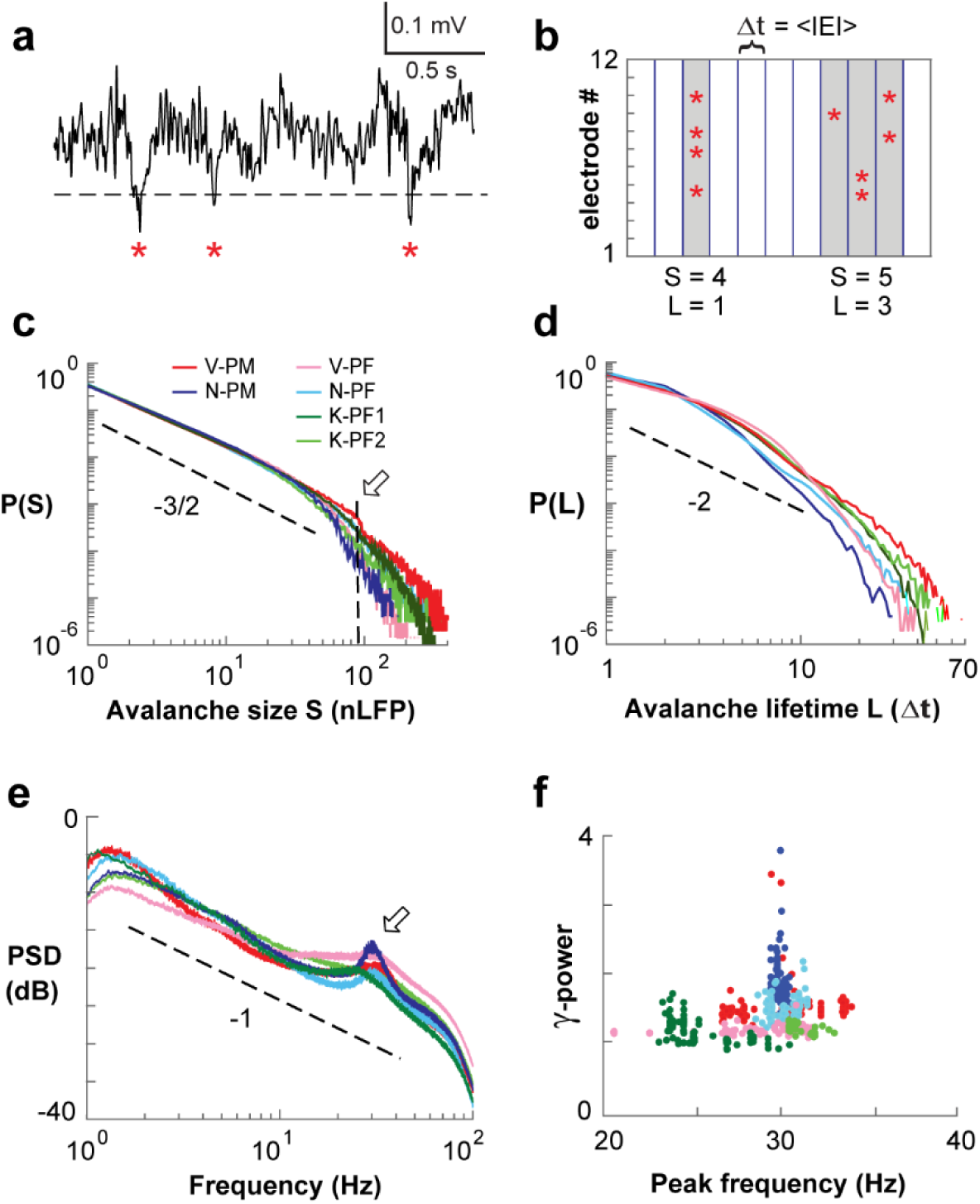
Neuronal avalanches and γ–oscillations coexist during resting activity in prefrontal (*PF*) and premotor (*PM*) cortex of nonhuman primates. **a**, Ongoing single electrode LFP with negative threshold crossing (2 s.d.; *dashed line*) defines nLFP times and peaks (*asterisks*). **b**, Concatenating successive time bins with at least 1 nLFP defines nLFP clusters of size 4, 5 (*gray*) at temporal resolution Δt on a 12-electrode array (*schematic*). **c**, Power law in nLFP cluster sizes identifies avalanche dynamics (Δt = ⟨IEI⟩). *Dashed line*: power law with slope of −3/2. *Arrow*: cut-off at ∼90 electrodes^55^. **d**, Non-power law duration distributions of nLFP clusters at Δt = ⟨IEI⟩. **e**, Power spectrum density (PSD) of the LFP with γ–activity peaks at 25–35 Hz. Average PSD per array (*color coded*) from single electrode PSDs. *Dashed line*: 1/*f*. **f**, Peak frequency vs. 1/*f*-corrected γ–power for each electrode. Legend in **c** applies to **d–f**.

### Exploration in temporal resolution identifies power law regimes for size and duration yielding **χ**_slope_ close to 2

Provided that both avalanche duration and size can be described by power laws, theory links the average size of avalanches with duration T, ⟨S⟩(T), to the avalanche profile^33, 34^. Specifically, the slope of ⟨S⟩(T) vs. T predicts the scaling exponent of avalanche profiles, χ_slope_. Yet, at Δt = ⟨IEI⟩, durations did not distribute according to a power law (see Fig. 1d), preventing an estimate of χ_slope_. Previous work has shown that an increase in Δt retains the power law in avalanche size while systematically reducing the corresponding slope α (Ref.^3, 12, 55–57^). Here we extended this approach to avalanche duration in the monkey to identify temporal resolutions for which both size and duration distributions of avalanches distribute according to power laws, potentially minimizing interference from γ–oscillations and thus allowing for an estimate of χ_slope_.

We assessed avalanche parameters for temporal resolutions Δt = 1–30 ms in steps of 0.5 ms and found that an increase in Δt systematically changed distributions for size and duration (Figs. 2a, b; see also Supplementary Fig. 1). Power laws were maintained for size (Fig. 2c; LLR >> 10^3^; vs. exponential; *P* << 0.005), with a steady shift in the cut-off and decrease in α (Fig. 2a, e). Importantly, whereas duration distributions failed the power law test for intermediate Δt = 2– 15 ms (Figs. 2d, f; *gray area*), they exhibited distinct power laws at higher and lower temporal resolutions (LLR >> 10^3^; vs. exponential; *P* < 0.005). Similar results were obtained when testing against log-normal alternative distributions (Supplementary Fig. 2). In the corresponding plots of ⟨S⟩(T) vs. T as a function of Δt, χ_slope_ was close to 2 for small and large Δt (Fig. 2g; *dashed lines*). This was quantified for all arrays by estimating χ_slope_ using linear regression based on lifetimes L ranging from n = 1–5, which avoids the cut-off in the lifetime distributions. For all arrays, χ_slope_ was found to be closer to 2 for valid power law duration regimes, whereas it approached 1–1.5 for intermediate temporal resolutions where lifetime distributions did not conform to power laws anymore (Fig. 2h; *c.f.* asterisks Fig. 2b, g).

**Fig. 2:**
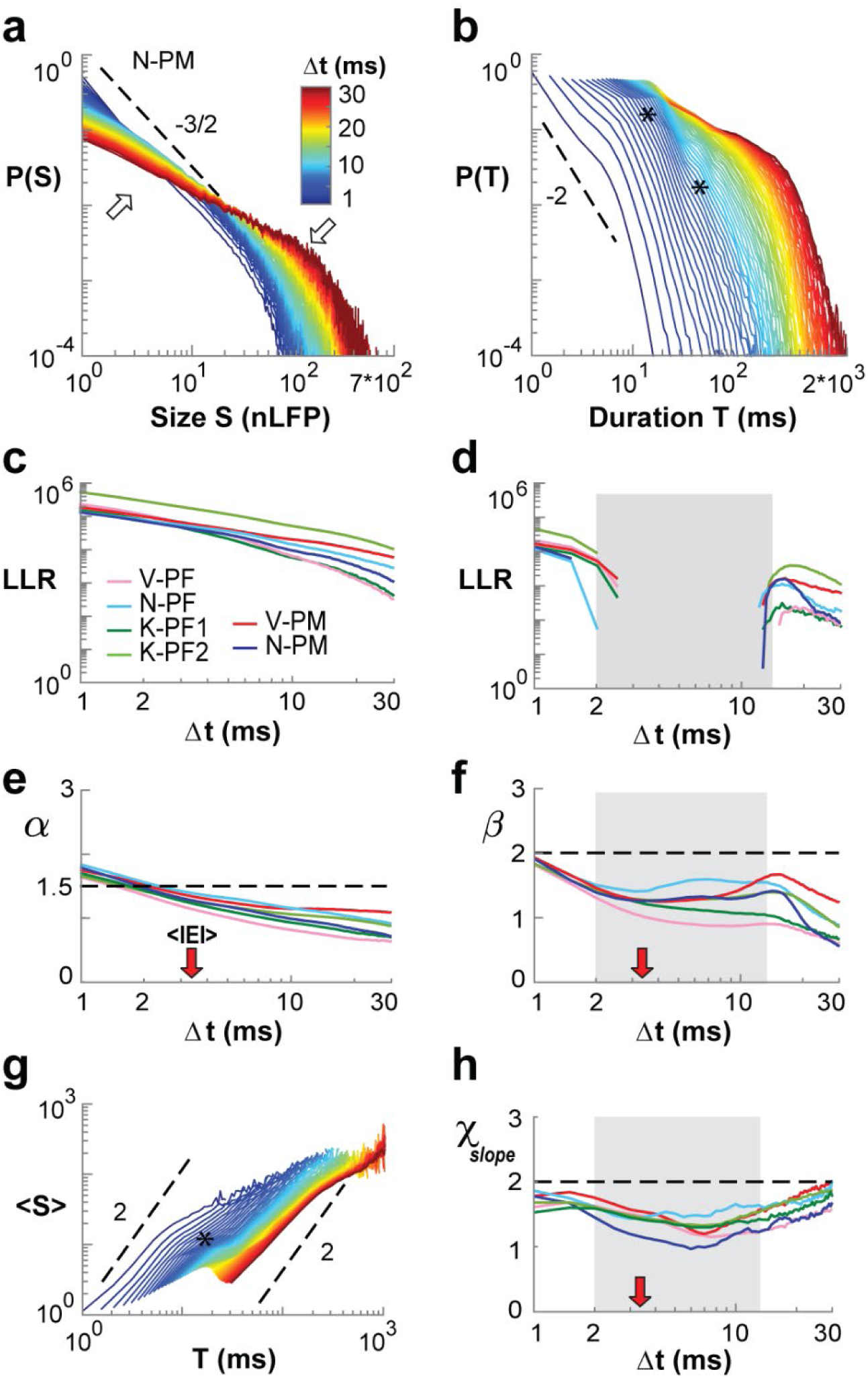
Systematic variation of temporal resolution yields power law regimes with χ_slope_ close to 2. **a**, Size distributions (single nonhuman primate; N-PM) decrease in slope (*upward arrow*) with increase in Δt and corresponding rightward shift in cut-off (*downward arrow*). **b**, Corresponding dependence of duration distribution on Δt. Note deformations (*asterisks*) at intermediate Δt. **c**, Size distributions remain power laws for all Δt (LLR values > 0 significant; vs. exponential distribution; all arrays; *color coded*). **d**, Corresponding summary for duration distributions demonstrating loss of power law for intermediate Δt (LLR < 0; *grey area;* all arrays). **e**, Summary of decrease in slope α as a function of Δt. **f**, Summary of slope β, which plateaus at intermediate Δt (*grey area taken from* d). **g**, Mean size-per-duration, ⟨S⟩(T), reveals deformations (*asterisks*) at intermediate Δt. **h**, χ_slope_ estimates are depressed at intermediate Δt. *Red arrows*: <IEI> for all arrays. *Dashed lines*: power law with given slope as visual guides. Legend in **a** applies to **b, g**. Legend in **c** applies to **d–f, h**.

We note that estimates of χ_slope_ for temporal resolutions more coarse than 30 ms revealed the increasing influence of the cut-offs in both size and lifetime distribution, demonstrated by n = 3 arrays that provided a sufficient number of avalanches under these conditions. When lowering the temporal resolution to 30, 50, 80 and 100 ms (Fig. 3), a rapid transition to the cut-off regime, beyond which avalanches cannot be accurately assessed^55^, dominates ⟨S⟩(T) with a corresponding change to χ_slope_ = 1. At this value, average avalanche size grows linearly with duration, which identifies an uncorrelated data regime comparable to shuffled data for which temporal correlations have been removed (Fig. 3; *colored dashed lines*). We conclude that the slope relationship for avalanche size and duration identifies a scaling exponent close to 2 at temporal resolutions that support power law regimes.

**Fig. 3:**
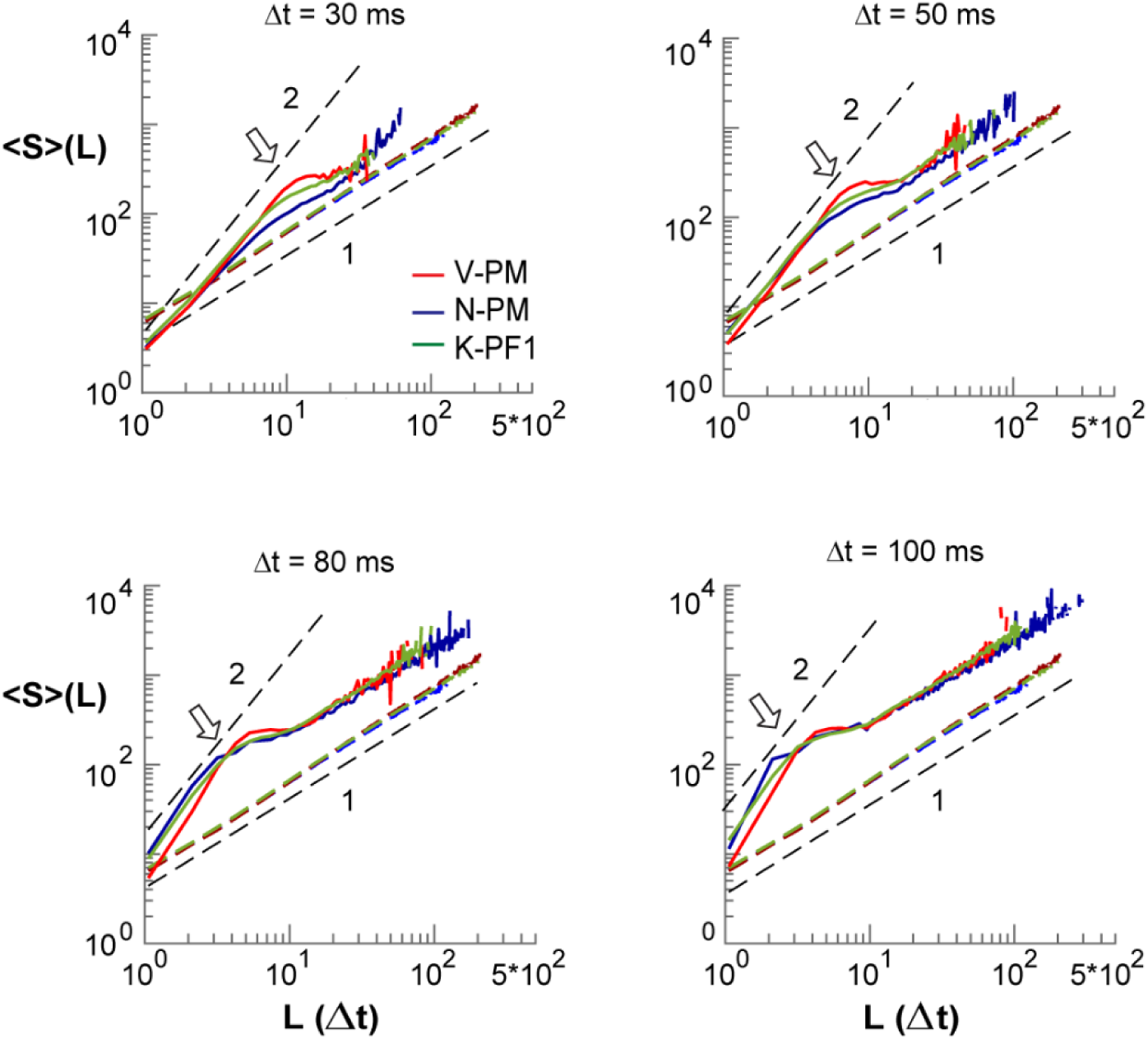
Shift in ⟨S⟩(L) power law-cutoff shows creeping transition from critical to linear trend for overly coarse temporal resolutions. Mean size-per-lifetime, ⟨S⟩(L), at time resolutions Δt = 30, 50, 80, 100 ms in a subset of arrays (*color coded*). *Black dashed lines*: visual guides for given power law slopes. *Arrows*: elbows in ⟨S⟩(L) are associated with the transition to P(S) and P(L) power law cutoffs, where cluster size simply grows linearly with cluster duration. *Black dashed lines*: visual guides for χ_slope_ = 2 and χ_slope_ = 1 respectively. *Colored dashed lines*: Corresponding ⟨S⟩(L) relationship for phase-shuffled data with χ_slope_ = 1 for uncorrelated temporal activity.

### The inverted parabolic profile of neuronal avalanches is modulated by γ–activity at intermediate Δt

To detail the impact of γ–activity on profile measures, we directly calculated avalanche profiles for different Δt. Critical theory predicts the avalanche profile to be an inverted parabola independent of duration T (Refs.^33, 34, 58^). Indeed, avalanche profiles followed an inverted-parabola at small and large Δt (Fig. 4a). The corresponding underlying continuous LFP time course revealed that at small Δt, profiles fit within a single γ–cycle, whereas profiles did not resolve γ–activity at large Δt. On the other hand, profiles deviated from a parabola at intermediate Δt for which they tracked multiple γ–cycles on the array (Fig. 4a; Δt = 3.5 ms, 10 ms). Accordingly, when analyzing avalanches of different lifetimes L, we find scale-invariant profiles at small and large Δt, but not at intermediate Δt (Fig. 4b; Δt = 9 ms, 15 ms). We quantified this relationship for all 6 arrays in corresponding error density plots in the (L, Δt)– plane. Strongest systematic deviations from a parabola were uncovered for T = L×Δt ≅ 66 ms, which spans ∼2 γ–cycles at 30 Hz (Fig. 4c; *dashed lines*). In contrast, error density plots based on a semicircle revealed reasonable fits mainly in the cut-off regime of avalanche duration distributions (Supplementary Fig. 3a).

**Fig. 4:**
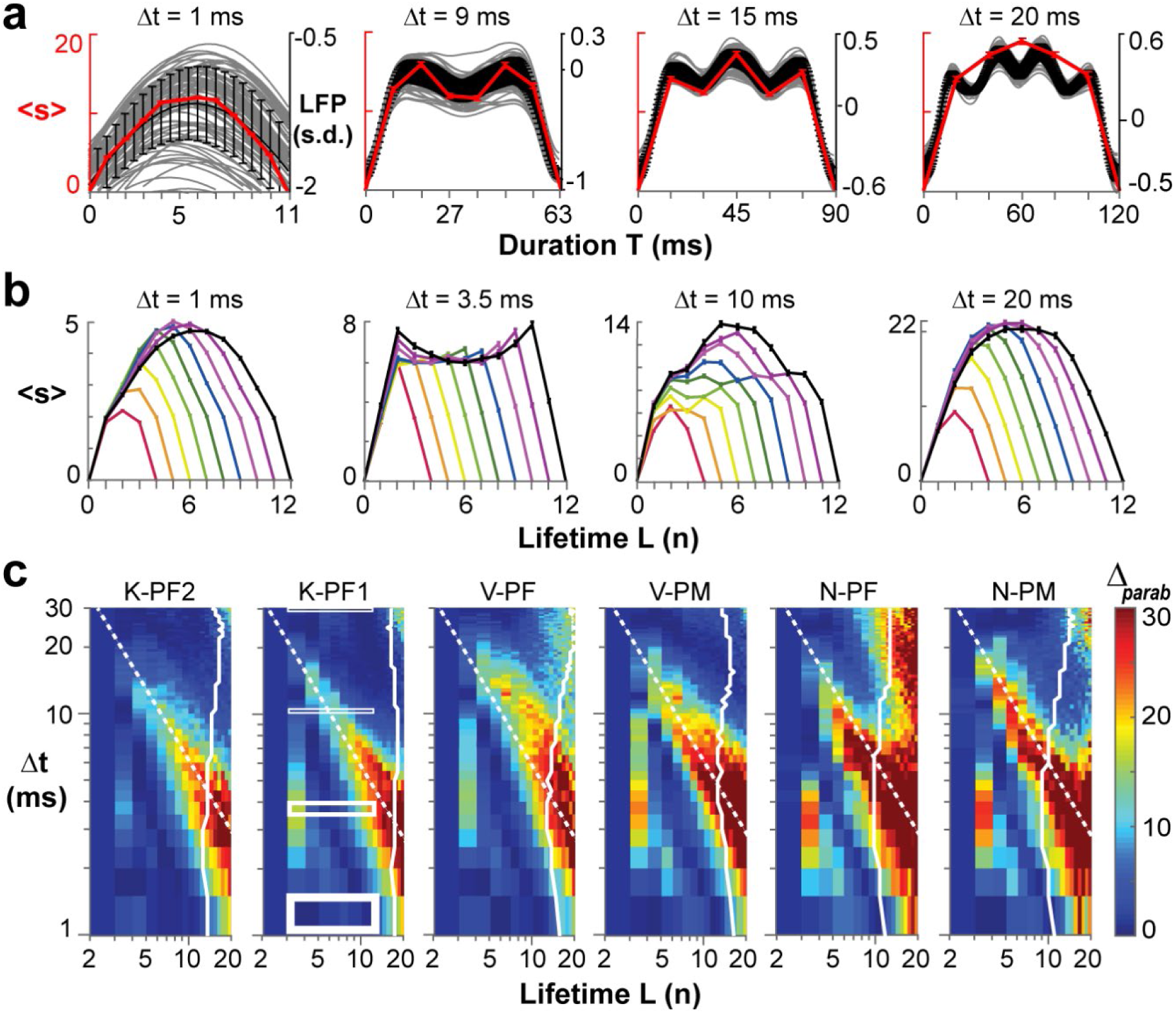
The inverted parabolic profile of neuronal avalanches is modulated by γ–activity at intermediate Δt. **a**, Examples of temporal avalanche profiles (*red*: mean size-per-timestep ± s.e.; N-PM) at increasing Δt and corresponding underlying γ–oscillations in the LFP (*grey*: mean inverted LFP of single electrode; *black*: mean ± s.d. of the array). **b**, Variable profiles are bracketed by parabolic profiles at small and large Δt for lifetimes L = 3, …, 11 (mean ± s.e.; K-PF1). **c**, Density plot of fit quality to an inverted parabola for all arrays and all profiles in the (L, Δt)–plane. Note compact regions of high deviations from a parabola surrounded by good parabolic fits. Rectangular regions in K-PF1 indicate profile ranges displayed in **b**. *Dashed lines*: high Δ_parab_ for condition T = L×Δt ≅ 66 ms.

### The scale-invariant, parabolic profile of neuronal avalanches reveals a scaling exponent **χ**_collapse_ **≅** 2

Critical theory predicts an exponent χ larger than 1.5 for the successful collapse of neuronal avalanche profiles into a single scale-invariant temporal motif 𝓕(t/L)^31–34^. Our demonstration of regimes with good parabolic fit (Fig. 4c) allowed us to identify 𝓕(t/L) directly over a range of L (Methods; Equation 1) by finding χ_collapse_, the estimate of χ that provided us with the lowest collapse error (Δ_F_) over an extended range of L and different Δt (Fig. 5). Example profiles for L = 3 to 11, their corresponding best global collapse, χ_collapse_, as well as residual profile differences together demonstrate good collapse with χ_collapse_ ≅ 2 at small and large Δt, but not at intermediate Δt (Fig. 5a–c). These findings on χ_collapse_ are in line with our findings on χ_slope_ derived from the ⟨S⟩(T) vs. T relationship. To further delineate the impact of γ–activity on profile collapse, we systematically identified non-collapsible regions in the (L, Δt)–plane by plotting Δ_F_ for the local collapse of (L–1, L, L+1) as a function of Δt (Fig. 5d). This approach revealed a band of high Δ_F_ for T = L×Δt ≅ 40 ms for all arrays, located at the transition from single- to double-peaked profiles. We note that the band of high Δ_F_ was solidly placed within the power law regime (Figs. 5d, *white solid line* indicates crossover to fewer than 1000 avalanches) and thus, does not indicate weak statistics due to e.g. a low number of avalanches as found beyond the cut-off (*c.f.* Figs. 1, 2). Use of a global collapse approach, i.e. L = 3, …, 20 revealed similar regimes of perturbation from γ–activity, yet with less resolution (Supplementary Fig. 3b).

**Fig. 5:**
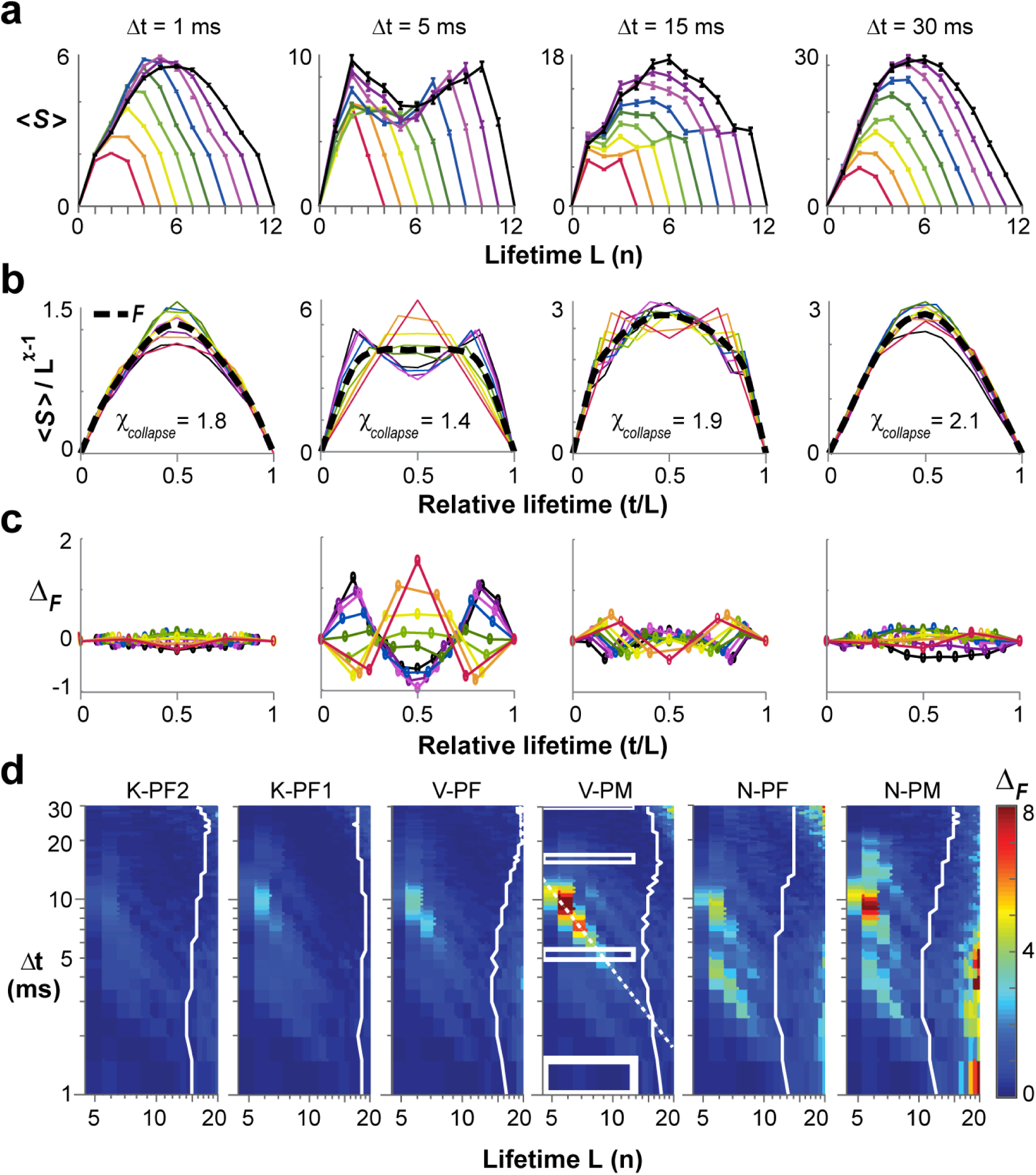
Scaling collapse of the inverted parabolic profile of neuronal avalanches. **a**, Examples of the average temporal profile ⟨S⟩(t, L) for avalanches of lifetime L = 3, …, 11 (*color coded*; mean ± s.e.) at temporal resolution Δt = 1, 5, 15 and 30 ms (single nonhuman primate; V-PM). **b**, Corresponding profile collapse with duration normalized by L and ⟨S⟩ scaled by *L*^χ−1^. χ_collapse_ obtained from best parabolic collapse 𝓕(t/L) (*solid black line*; see Methods). **c**, Profile collapse differences, Δ_F_, obtained by subtracting best collapse 𝓕(t/L) from normalized profiles are largely symmetrical around profile peak. **d**, Density plot of collapse error for all 6 arrays ranked from low (*left*) to high γ–oscillation power (*right*). Collapse error obtained from profiles L−1, L, and L+1 for given Δt. Collapse failure visible for intermediate Δt before the cut-off in lifetime distribution (*white border* equals threshold for L with <1000 avalanches indicating a statistical cut-off regime).

If γ–activity was a major factor in preventing proper avalanche profile collapse, we expect the collapse error to correlate with the strength of γ–activity. Indeed, the average collapse error, ⟨Δ_F_⟩, significantly correlated with peak γ–power across arrays, confirming that the band of high Δ_F_ reflects the impact of γ–oscillations (Fig. 6a; *P* = 0.0054). For temporal resolutions of 1 ms and 30 ms, which were found to be least affected by γ–activity, the corresponding collapsed motif 𝓕 was significantly closer to a parabola than a semicircle (Fig. 6b; Δ_parab_ < Δ_semi_, ANOVA, *P* = 1.4×10^−6^; *F* = 22.2), the latter being predictive of non-critical dynamics^38^. The finding of χ_collapse_ ≅ 2 for these temporal resolutions and the deviations from 2 for intermediate resolutions is summarized for all arrays in Fig. 6c.

**Fig. 6:**
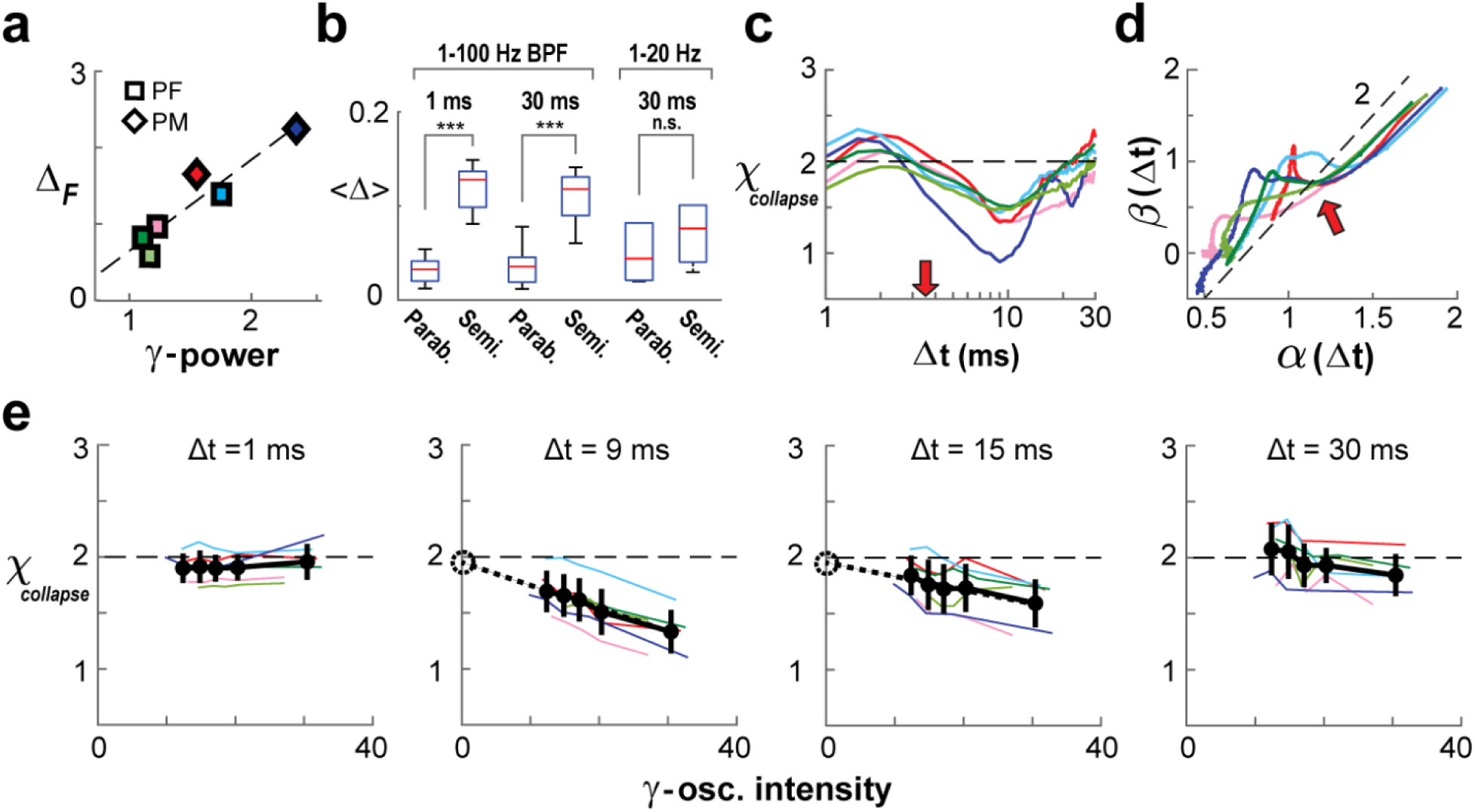
Derivation of scaling exponent χ_collapse_ ≅ 2 for the parabolic temporal profile of neuronal avalanches in the presence of γ–oscillations. **a**, Average collapse error positively correlates with peak γ–power (based on PSD; *c.f.* Fig. 1e, f) across arrays (*P* = 0.005). **b**, A parabola consistently fits the temporal avalanche profile better than a semicircle for high and low temporal resolutions (*1–100 Hz BPF*). However, when γ–oscillations were filtered out (*1–20 Hz*), a semicircle fit could not be rejected. Error bars represent array mean ± s.d. **c**, Summary of χ_collapse_ obtained as a function of Δt (temporal profile collapse L = 3, 4, 5). χ_collapse_ is close to 2 outside γ–activity impact. **d**, Estimation of scaling exponent χ_ratio_ follows a slope of ≅ 2 when plotting slope estimate β vs. α as a function of Δt. Note strong deviation for intermediate Δt indicating impact of γ–oscillations. *Dashed line*: slope of 2 as a visual guide. *Red arrow*: <IEI>. **e**, Array-averages of χ_collapse_ for five quintiles of γ–intensity (black lines indicate mean ± s.d. across arrays; see Methods) reveal agreement with the expected value of 2 (*dashed lines*) for high and low temporal resolutions. *Colored lines*: individual arrays. Intermediate resolutions of 9 and 15 ms consistently show χ_collapse_ < 2 with largest deviation seen in high wave-impact quintiles; however, a linear fit of the average trend (*dotted lines*) intersects with the y-axis at approximately χ_collapse_ = 2 (*dashed circle*). For *color code c.f.* Fig. 1c.

Theory^34^ predicts that the quotient of size and duration slopes, (β−1)/(α−1), provides the scaling exponent of avalanche profiles, χ_ratio_. Although individual slope estimates are a function of Δt, we asked whether the ratio of slope estimates might support a critical scaling exponent of χ_ratio_ ≅ 2. Accordingly, when plotting β(Δt) vs. α(Δt) we find that χ_ratio_ follows closely a slope of ≅ 2 except for intermediate Δt (Fig. 6d).

So far, we have shown that straightforward estimates of χ are not possible at temporal resolutions that appropriately resolve γ–activity. Next, we devised an approach that allowed us to estimate the value of χ_collapse_ in the limit of vanishing γ–activity for temporal resolutions that accurately track multiple γ–cycles. This approach takes advantage of spontaneous moment-to-moment fluctuations in ongoing γ–activity. We first sorted subsequent 1-second intervals of ongoing activity into quintiles according to their absolute γ–intensity (see Methods) and then obtained a collapse in avalanche profiles for each quintile (Fig. 6e). In line with expectations, we obtained estimates of χ_collapse_ ≅ 2 that were independent of γ–intensity across quintiles for Δt = 1 ms and Δt = 30 ms. Furthermore, as expected for intermediate temporal resolutions, χ_collapse_ depended on γ–intensity, being larger for quintiles with less γ–intensity. Importantly, regression estimates of χ_collapse_ in the limit of no γ–influence predicted χ_collapse_ ≅ 2 (Fig. 6e, *dotted lines*; Δt = 9ms, χ_collapse_ = 1.95 **±** 0.03, slope −0.02, *P* < 0.001; Δt = 15 ms, χ_collapse_ = 1.96 **±** 0.04, slope −0.01, *P* = 0.01; linear regression, mean ± s.e.). In summary, our analysis robustly supports a critical scaling exponent of χ ≅ 2 for the collapse of avalanche profiles based on three estimates of this exponent: χ_slope_, χ_ratio_ and χ_collapse_.

### Filtering γ–activity removes critical signatures in the avalanche profile

If γ–activity was simply linearly superimposed on an otherwise scale-invariant avalanche background, then bandpass filtering between 1–20 Hz should reduce γ–activity without significantly changing estimates of avalanche parameter. We found that bandpass filtering reduced the rate of nLFPs by a factor of 4 to 5, yet a sufficient number of avalanches were obtained for n = 3 arrays (⟨IEI⟩ = 15.2 ± 1.6 ms; *n* = 3; Fig. 7). To account for the longer ⟨IEI⟩, we extended our analysis for Δt up to 100 ms. Power laws for size distributions were maintained for all temporal resolutions (LLR >> 10^3^; *P* < 0.005). However, duration distributions now failed the power law test for a new range of Δt = 8–25 ms, which was closer to the scale of β-oscillations (Fig. 7a, b; grey area; Supplementary Fig. 4) and for which χ_slope_ was close to 1.5 (Fig. 7b; χ_slope_). Similarly, profile collapses previously affected at the scale of γ–activity improved, (Fig. 7d, e; Δt = 5 ms), but double-peaked profiles shifted to lower resolutions with profile collapse strongly affected for T = L×Δt ≅ 75 ms, equivalent to the transition from one to two β–cycles at 20 Hz (Fig. 7c–f; Δt = 15 ms; *dashed line*). Importantly, profile collapse at Δt = 1 and Δt = 30 ms now revealed χ_collapse_ ≅ 1.5 (Fig. 7d; *c.f.* χ_slope_ in Fig. 7b), and profiles were not significantly different from a semicircle (Fig. 6b; 1–20 Hz). Our findings demonstrate that spontaneously arising γ–activity is intrinsically embedded within critical avalanche dynamics.

**Fig. 7:**
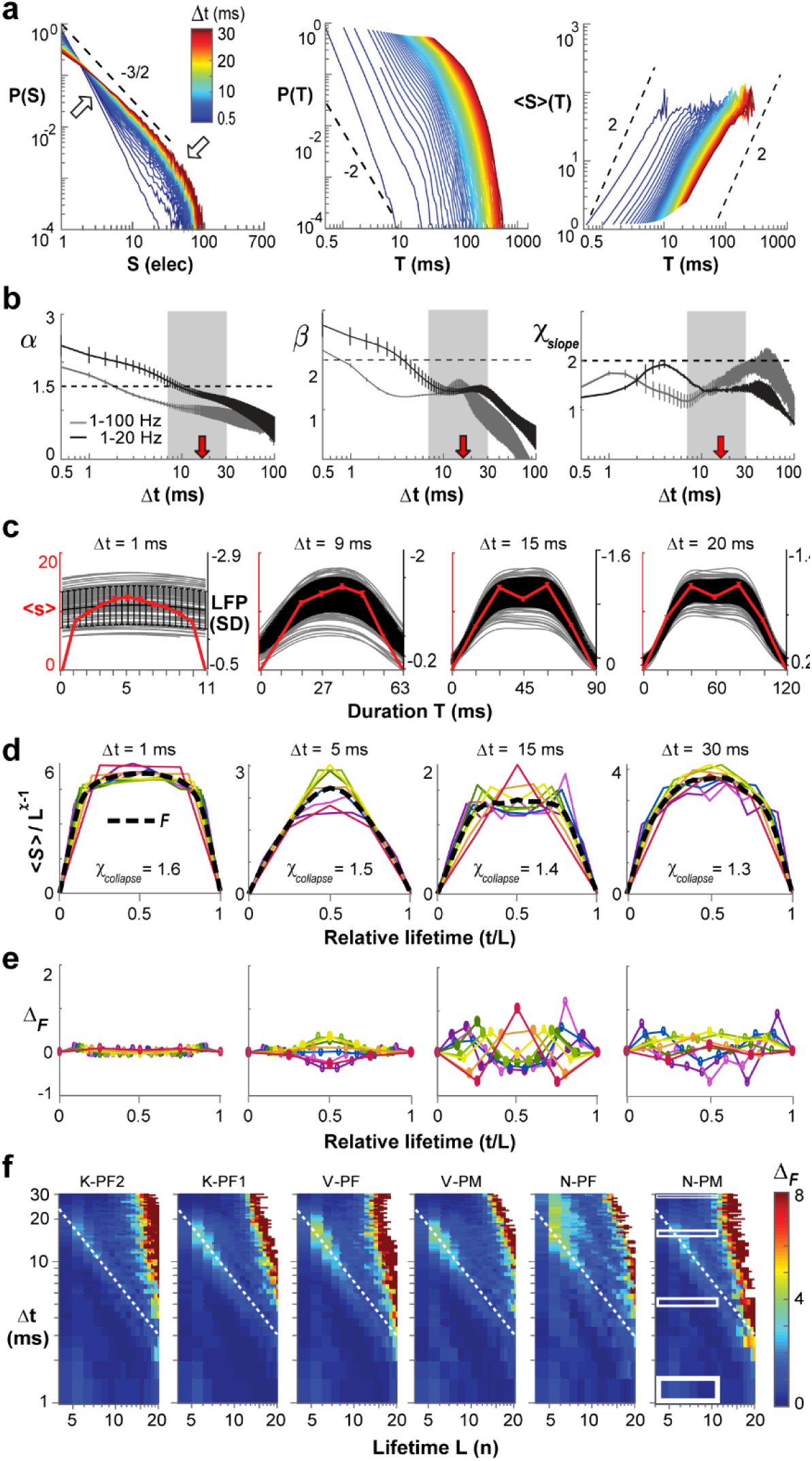
Filtering of γ–activity reduces χ_slope_ and χ_collapse_ and changes the temporal profile of neuronal avalanches to a semicircle. **a**, Avalanche size and durations distributions, as well as mean size-per-duration ⟨S⟩(T) as a function of Δt for data bandpass filtered between 1–20 Hz (Butterworth 6^th^ order). Note change in χ_slope_ towards values smaller than 2 in the ⟨S⟩(T) plot (*c.f.* Fig. 2h). **b**, Smooth decrease in α with increasing Δt (*left*), yet variable slope changes for β (*middle*) and χ_slope_ (*right*) at intermediate Δt. *Grey*: 1–100 Hz mean ± s.d. taken from Fig. 1 for comparison; average over 3 monkeys. *Black*: corresponding 1–20 Hz mean ± s.d.. *Red arrows:* ⟨IEI⟩ for 1–20 Hz. *Black dashed lines*: visual guides of theoretical power law slopes. *Grey area*: loss of power law in duration distributions for intermediate Δt (LLR < 0; all arrays). **c**, Examples of continuous LFP waveforms and corresponding avalanche profiles. **d**, Collapse in temporal profiles over a wide range of temporal resolutions Δt (*c.f.* Fig. 4, here L = 1, …, 9 color coded from red to purple). Best collapse is obtained for χ_collapse_ close to 1.5 and lower. **e**, Symmetrical difference of normalized profiles from best fit (*c.f.* Fig. 5c). **f**, Density plot of collapse error Δ_F_ obtained from profiles L−1, L, and L+1 for given Δt. *Dotted lines*: high collapse error Δ*_F_* for condition T = L×Δt ≈ 80 – 100 ms, which equals about two β-cycles. Rectangular regions in N-PM indicate profile ranges displayed in a, c, d, and e.

## DISCUSSION

Identifying constraints in how neuronal activity changes in time is central to understanding cortex function. The most commonly reported temporal organization is that of an oscillation, found in cortical activity at separate physiological frequency ranges such as the theta (θ; ∼4–8 Hz), beta (β; ∼12–30 Hz) and gamma bands (γ; ∼30–100 Hz)^44–46, 59^. The oscillation phase provides a distinct scale for neuronal synchronization that is central to many theories on cortical processing^47, 60–62^. In contrast, very different dynamics that also capture neuronal synchronization have been found in the form of neuronal avalanches^3^. The present work uncovers a scale-invariant temporal parabola in the organization of ongoing neuronal avalanches, a finding in line with expectation for critical dynamics. Previous work in *in vitro* and in rodents reported nested θ/β/γ–oscillations^63, 64^ embedded in avalanches. Simulations also demonstrate that oscillations can emerge during avalanches^65–68^. However, it had yet to be determined whether cortical avalanches in the awake mammal exhibit a distinct temporal profile that is scale-invariant, nor had it been known how that profile might relate to the scale of an oscillation. Here, we demonstrate the motif of an inverted parabola that guides the scale-invariant temporal evolution of neuronal avalanches in the propagation of ongoing cortical activity *in vivo*. This motif uncovers a specific constraint for ongoing cortical dynamics, that is, local events initiate spatial expansion that is similar at all scales and collapses symmetrically in time. Thus, despite the complex temporal evolution of activity within an avalanche, the average profile exhibits time reversal^32^.

This motif and its corresponding collapse are predicted by the theory of critical systems^31–34, 58^, which also states that the critical exponents of the power law distribution in size, α, and duration, β, relate to the collapse exponent as 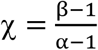 (Ref.^34^). For neuronal avalanches, empirical^3^ and simulated^69^ slopes of α are close to an exponent of 3/2 at Δt = ⟨IEI⟩. These empirical insights are supported within the Landau-Ginzburg theoretical framework of network dynamics^50^. The size exponent of 3/2 is also characteristic for a critical branching process in line with the well-established empirical finding of a critical branching parameter of 1 for neuronal avalanches^3^. Critical branching processes exhibit a lifetime exponent β = 2, leading to a corresponding χ = 2. However, empirically obtained lifetime distributions have been difficult to analyze due to their typically narrow range in lifetimes^3, 8^. Our analysis identifies a new element affecting lifetime distributions by showing that the exponent β cannot be reliably determined at ⟨IEI⟩ in the presence of γ–oscillations, leading to a corresponding depression in χ. Importantly, when the impact of the oscillation is adequately taken into account, χ comes close to 2, as expected for critical avalanche dynamics. This in turn supports a critical exponent for avalanche lifetimes of β = 2. Our demonstration of χ = 2 in the awake monkey based on LFPs represents a significant departure from estimates of χ = 1.2 – 1.5 based on spike activity^70^, which suffer from subsampling^71^.

Scaling collapse of avalanche profiles has been increasingly used to study avalanche dynamics beyond the identification of power laws in size and duration (e.g Ref^35^). While profiles utilize more information about avalanches than size or lifetime distributions, robust profile collapse requires orders of magnitude more data compared to distribution estimates. Our current analysis was based on ∼3 million nLFP events per array (see Supplementary Tables 1 & 2) with a correspondingly high number of avalanches allowing for the reconstruction of robust profiles even for long lifetimes, which are rare (>7Δt; *c.f.* Fig. 1). This in turn enabled us to identify temporal resolutions of the parabolic motif with high collapse quality and also allowed us to distinguish the modulating influence of γ–oscillations from profile variations due to reduced statistical power. We note that, by definition, avalanches start with the peak time of an nLFP, which synchronizes avalanches to the first oscillation peak on the array.

We demonstrated that removal of γ–oscillations by filtering abolished the critical signature of the temporal avalanche profile. The resulting profile was similar to the semicircle motif with a scaling exponent of *χ* ≅ 1.5 reported for non-critical cascading models (see e.g. Ref.^38^). Following filtration of fast components, the variability in avalanche profile and corresponding changes in slope estimates were consequently found to shift to the scale of β–oscillations. We interpret these findings to mean that at the spatial scale of the microelectrode array (∼400 μm interelectrode distance) and the temporal scale of γ–oscillations (5–10 ms), γ– activity needs to be considered to fully capture the critical aspect of neuronal avalanches. Substantial coarse graining in space and time might be required to identify the contribution of lower frequency oscillations such as low-β and θ oscillations in order to identify potential contribution of those frequency bands to the temporal profile of avalanches. We found that slope estimates for temporal resolutions coarser than 30 ms (Fig. 3) were dominated by uncorrelated activity. This suggests that experimental techniques different from LFP recordings with high-density microelectrode arrays might be required to identify the interdependence between critical dynamics and oscillations lower than 20 Hz. Nevertheless, our approach presented here might guide the analysis of avalanches found at lower temporal and spatial scales, for example, in human fMRI^19^. We note that scaling collapses have also been successfully employed to quantify the temporal organization of behavior^72, 73^ at time scales of minutes to hours.

Our identification of a symmetrical (in time), parabolic profile for neuronal avalanches *in vivo* supports results from simulations of critical neural networks^35, 56^ and identifies constraints for other generative models proposed for avalanche dynamics^37^. Models fine-tuned to produce power laws for size and duration distributions via bi-stable dynamics combined with a locally expansive dynamical term^39^ or purely external uncorrelated driving^38^ reveal profiles that deviate from a symmetrical parabola. The parabolic profile also differentiates neuronal avalanches from stochastic processes with no memory, which typically display a semicircle motif^74^. Specific graph-theoretical constructs such as a hierarchical topology^40^ can also mimic scale-free size distributions but fail to produce an inverted parabola profile. Network and biophysical models have demonstrated avalanches to emerge with oscillations at a particular E/I balance or topology; however, the corresponding avalanche profile was not reported^66, 75, 76^. Our findings are in line with models showing the coexistence of avalanches and neuronal oscillations at a continuous synchronization/desynchronization phase transition^50, 68^. These models do not include transmission delays but exhibit exponents expected for a critical branching process. Less abstract analysis regarding the coexistence of critical and oscillatory dynamics must incorporate biologically relevant transmission delays to which neuronal oscillations are known to exhibit high sensitivity^77, 78^.

It is well accepted that limited predictability is one of the major disadvantages of critical systems^79^. Our findings support recent suggestions^80^ that the coexistence of oscillations with critical dynamics might allow the brain to combine functional benefits of criticality for information processing^23–30, 81^ with temporal precision. Such precision might be important for behavioral outcomes or learning that have been typically studied in isolation from underlying scale-free fluctuations^47–49^.

The precise profile of neuronal avalanches might be a suitable biomarker for pathological brain dynamics. In human preterm infants suffering from anoxia, asymmetric avalanche profiles in ongoing EEG activity became symmetrical upon recovery^41, 42^, which is supported by our demonstration of a symmetrical profile during normal activity. Furthermore, γ–activity has been identified as a key marker in disease states such as schizophrenia^82^. Our demonstration that γ– activity and avalanche profiles are non-linearly related supports the idea that mental disorders such as schizophrenia might indicate deviations from critical brain dynamics^83^.

## METHODS

### Animal procedure

All animal procedures were conducted in accordance with NIH guidelines and were approved by the Animal Care and User Committee of the National Institute of Mental Health (Animal Study Protocol LSN-11). Three adult nonhuman primates (*Macaca mulatta*; 1 male, 2 females; 7–8 years old) received two chronic implantations of high-density 96-microelectrode arrays (Blackrock Microsystems; 4×4 mm^2^; 400 µm interelectrode distance; 10×10 grid with corner grounds). To direct recordings towards superficial cortical layer II/III, electrode shanks of 0.6 mm length were used in prefrontal cortex (PF; *n* = 4), and shanks of 1 mm length were used in premotor cortex (PM; *n* = 2). During recording sessions, monkeys sat head-fixed and alert in a monkey chair with no behavioral task given. Portions of this dataset have been analyzed previously^55, 84^.

### Electrophysiological recordings and preprocessing

Simultaneous and continuous extracellular recordings were obtained for 12–60 min per recording session (2 kHz sampling frequency), filtered between 1–100 Hz (6^th^-order Butterworth filter) to obtain the LFP and notch-filtered (60 Hz) to reduce line noise. Arrays on average contained 85 ± 8 functional electrodes that exhibited 64 ± 50 µV of spontaneous LFP fluctuations (s.d.). About 7 ± 3% of time periods were removed from functional electrodes due to artifacts introduced by e.g. vocalization, chewing, sudden movements. These artifacts were identified by threshold crossing (s.d. > 7) and excised (±0.25 s). Electrode LFPs were z-transformed and recording sessions for each array were combined for further analysis. The current study represents a combined 22 hr of ongoing cortical LFP activity (for details see Supplementary Table 1).

### Power spectrum analysis

Single electrode periodograms (Mathworks; *pwelch*) were averaged for each array and displayed in double-logarithmic coordinates. The power of prominent γ– oscillations was quantified for each electrode by subtracting a full-spectrum power-matched 1/*f* fit from the original periodogram and dividing the area under the resulting curve from *f*_peak_ ± 1 Hz by the integral over the range *f*_peak_ ± 5 Hz.

### Avalanche definition

For each electrode in the array, absolute peak amplitude and time of negative LFP threshold crossings (–2 s.d.) were extracted at Δt = 0.5 ms and combined into a matrix, i.e. raster, with rows representing electrodes and columns representing time steps. Continuous amplitude rasters were converted to binary rasters by setting amplitude values to 1 and rasters at reduced temporal resolutions were obtained by concatenating columns. A population time vector was obtained by summing nLFPs in the raster for each time step, and avalanches were defined as spatiotemporal continuous activity in the population vector bracketed by at least one time-bin with no activity. The size of an avalanche, S, was defined as the number of nLFPs participating. Multiple nLFPs at an electrode during an avalanche are rare^55^ and were counted in size estimates. Avalanche lifetime, L, was defined as the number of successive time bins, n, spanned by an avalanche in multiples of temporal resolution Δt. Avalanche duration, T, was expressed in absolute time as T = L×Δt. Scale invariance of S, L and T was visualized by plotting probability distributions P(S), P(L), and P(T) in log-log coordinates.

### Statistical tests and slope estimates

The maximum-likelihood ratio (LLR) was calculated to test potential power law distributions against the alternatives of exponential distributions if not stated otherwise^52^. As a control, we also tested against log-normal distributions. Significance of the LLR was determined according to previously published methods^51, 52^ and confirmed LLR results.

Estimations of initial slope were based on linear regression of log-converted data excluding higher cut-off effects. For α, we used the range of S = 1 to 40, which excludes the distribution cut-off close to the total number of functional electrodes on the array (*n* ≥ 70). Initial slopes β of P(L) and χ of ⟨S⟩(L) were estimated from L = 1–5.

The estimate for χ_slope_ for shuffled data was based on phase-shuffling the original time series at sampling frequency in the Fast Fourier domain followed by a retransformation and performing avalanche analysis on the reconstructed time series.

An ANOVA with Tukey’s post hoc correction was used to test for class differences if not stated otherwise.

### Collapse of the temporal profile of avalanches

Avalanches were grouped by L in multiples of Δt and averaged to obtain the average temporal profile for a given lifetime, ⟨S⟩(t, L). Normalized to dimensionless time units, t/L, amplitudes were then rescaled via Equation 1.

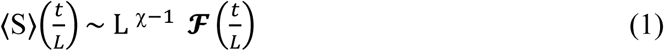

The profile collapse function, shown in Equation 1, relates the mean profile for each lifetime L, ⟨S⟩(t/L), with a characteristic temporal motif 𝓕(t/L), and scaling factor, L^χ−1^, which is independent of L according to Equation 1. To perform a shape collapse, we plotted ⟨S⟩(t, L) from L−1 through L+1 for L_min_ = 3Δt (to reduce finite size effects in shape caused by too few data points) through L = L_max_ chosen based on the strength of our data set (>1,000 avalanches per *L*). The collapse error, Δ_F_, was quantified via a normalized mean squared error (NMSE) of height-normalized individual profiles to the combined normalized average of all collapsed profiles, 𝓕(t/L). Minimized collapse error was calculated by scanning through χ = 0.5 to 3 at resolution of 0.001 to find the collapse in avalanche waveform associated with the smallest Δ_F_ via χ_collapse_. A value of Δ_F_ > 1 was considered a failure in collapse.

### Parabolic fit

For the parabolic fit we used the approach by Laurson^32^ as follows:

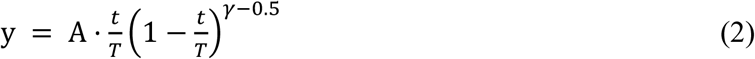

The parabolic fit error, Δ_parab_, was quantified via a normalized mean squared error (NMSE) of individual profiles to an amplitude-matched parabola which was coarse-grained to match L. Comparison to a semicircle fit was conducted in the same manner to obtain Δ_semi_ using:

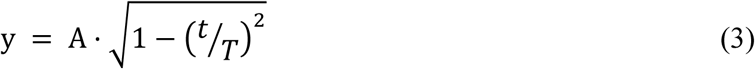

### LFP waveforms

To trace the oscillation origin of deviations from the parabolic shape motif, temporal profiles of neuronal avalanches were compared to their underlying LFP waveforms with sampling frequency of 2 kHz. Waveforms were grouped by L and averaged. Single electrode averages were used to calculate the array average.

### Limit estimate of χ_collapse_

To estimate χ_collapse_ at temporal resolutions affected by γ–oscillations, we classified ongoing 1 s segments of ongoing activity according to their γ–intensity. We first bandpass filtered each LFP recording from 20-40 Hz to isolate the presence of 30 Hz gamma bursts. Next, the Hilbert transform was taken and squared to acquire the corresponding γ– intensity time series for each electrode on the array. We then stepped through each recording in 1 second increments, labeling each segment with the maximum intensity observed at any electrode during that second. Time segments were regrouped as a function of peak γ–intensity into quintile classes. We then stitched together the new regrouped time series for each class across recordings and performed a full analysis on each quintile. The average intensity within each class was used to quantify the effect of γ–activity on the scaling exponent χ_collapse_, and results were averaged across monkeys and cortical areas.

## Acknowledgements

We thank Drs. Rajarshi Roy and Wolfgang Losert for their excellent mentoring of S.R.M., who was a Ph.D. student at the University of Maryland College Park Institute for Physical Sciences and Technology. We thank Drs. Lins-Ribeiro, Chris Meisel and Yahya Karimipanah as well as attendees of the Fifth Annual International Workshop on Criticality and the Brain for their support and useful discussions. This research utilized the computational resources of the NIH HPC Biowulf cluster (http://hpc.nih.gov), and was supported by the Intramural Research Program of the National Institute of Mental Health ZIA MH00297, USA.

## Author Contributions

S.Y. collected the data; S.R.M. analyzed the data; S.R.M. and D.P. wrote the manuscript.

## Supplementary Information

See Supplementary Info file.

### Competing interests

The authors declare no competing interests.

## Notes

#### Summary of Updates

Sections on Figs. 6 & 7 updated for clarification

## REFERENCES

1 Kreindler, G. E. & Young, H. P. Rapid innovation diffusion in social networks. Proc. Natl. Acad. Sci. U.S.A. 111, 10881–10888 (2014).

2 Pastor-Satorras, R., Castellano, C., Van Mieghem, P. & Vespignani, A. Epidemic processes in complex networks. Rev. Mod. Phys. 87, 925 (2015).

3 Beggs, J. M. & Plenz, D. Neuronal avalanches in neocortical circuits. J. Neurosci. 23, 11167–11177 (2003).

4 De Domenico, M., Granell, C., Porter, M. A. & Arenas, A. The physics of spreading processes in multilayer networks. Nat. Phys. 12, 901–906 (2016).

5 Alstott, J., Pajevic, S., Bullmore, E. & Plenz, D. Opening bottlenecks on weighted networks by local adaptation to cascade failures. J. Complex Netw. 3, 552–565 (2015).

6 Gleeson, J. P. & Durrett, R. Temporal profiles of avalanches on networks. Nat. Commun. 8, 1227 (2017).

7 Yu, S. et al. Higher-order interactions characterized in cortical activity. J. Neurosci. 31, 17514–17526 (2011).

8 Plenz, D. Neuronal avalanches and coherence potentials. Eur. Phys. J. Spec. Top. 205, 259–301 (2012).

9 Bellay, T., Klaus, A., Seshadri, S. & Plenz, D. Irregular spiking of pyramidal neurons organizes as scale-invariant neuronal avalanches in the awake state. eLife 4, 1–25 (2015).

10 Scott, G. et al. Voltage imaging of waking mouse cortex reveals emergence of critical neuronal dynamics. J. Neurosci. 34, 16611–16620 (2014).

11 Ponce-Alvarez, A., Jouary, A., Privat, M., Deco, G. & Sumbre, G. Whole-brain neuronal activity displays crackling noise dynamics. Neuron, 1446–1459 (2018).

12 Petermann, T. et al. Spontaneous cortical activity in awake monkeys composed of neuronal avalanches. Proc. Natl. Acad. Sci. U.S.A. 106, 15921–15926 (2009).

13 Yu, S. et al. Maintained avalanche dynamics during task-induced changes of neuronal activity in nonhuman primates. eLife 6, e27119 (2017).

14 Meisel, C. et al. Intrinsic excitability measures track antiepileptic drug action and uncover increasing/decreasing excitability over the wake/sleep cycle. Proc. Natl. Acad. Sci. U.S.A. 112, 14694–14699 (2015).

15 Solovey, G., Miller, K. J., Ojemann, J. G., Magnasco, M. O. & Cecchi, G. A. Self-regulated dynamical criticality in human ECoG. Front. Integr. Neurosci. 6 (2012).

16 Shriki, O. et al. Neuronal avalanches in the resting MEG of the human brain. J. Neurosci. 33, 7079–7090 (2013).

17 Palva, J. M. et al. Neuronal long-range temporal correlations and avalanche dynamics are correlated with behavioral scaling laws. Proc. Natl. Acad. Sci. U.S.A. 110, 3585–3590 (2013).

18 Arviv, O., Medvedovsky, M., Sheintuch, L., Goldstein, A. & Shriki, O. Deviations from critical dynamics in interictal epileptiform activity. J. Neurosci. 36, 12276–12292 (2016).

19 Tagliazucchi, E., Balenzuela, P., Fraiman, D. & Chialvo, D. R. Criticality in large-scale brain fMRI dynamics unveiled by a novel point process analysis. Front. Physiol. 3 (2012).

20 Cocchi, L., Gollo, L. L., Zalesky, A. & Breakspear, M. Criticality in the brain: A synthesis of neurobiology, models and cognition. Prog. Neurobiol. 158, 132–152 (2017).

21 Hesse, J. & Gross, T. Self-organized criticality as a fundamental property of neural systems. Front. Syst. Neurosci. 8 (2014).

22 Chialvo, D. R. Emergent complex neural dynamics. Nat. Phys. 6, 744–750 (2010).

23 de Arcangelis, L., Perrone-Capano, C. & Herrmann, H. J. Self-organized criticality model for brain plasticity. Phys Rev.Lett 96, 028107 (2006).

24 Kinouchi, O. & Copelli, M. Optimal dynamical range of excitable networks at criticality. Nat. Phys. 2 348–351 (2006).

25 West, B. J., Geneston, E. L. & Grigolini, P. Maximizing information exchange between complex networks. Phys. Rep. 468, 1–99 (2008).

26 Shew, W. L., Yang, H., Petermann, T., Roy, R. & Plenz, D. Neuronal avalanches imply maximum dynamic range in cortical networks at criticality. J. Neurosci. 29, 15595–15600 (2009).

27 Shew, W. L., Yang, H., Yu, S., Roy, R. & Plenz, D. Information capacity is maximized in balanced cortical networks with neuronal avalanches. J. Neurosci. 31, 55–63 (2011).

28 Yang, H., Shew, W. L., Roy, R. & Plenz, D. Maximal variability of phase synchrony in cortical networks with neuronal avalanches. J. Neurosci. 32, 1061–1072 (2012).

29 Gautam, H., Hoang, T. T., McClanahan, K., Grady, S. K. & Shew, W. L. Maximizing sensory dynamic range by tuning the cortical state to criticality. PLoS Comput. Biol. 11, e1004576 (2015).

30 Shriki, O. & Yellin, D. Optimal information representation and criticality in an adaptive sensory recurrent neuronal network. PLoS Comput. Biol. 12, e1004698 (2016).

31 Rybarsch, M. & Bornholdt, S. Avalanches in self-organized critical neural networks: A minimal model for the neural SOC universality class. PLoS One 9, e93090 (2014).

32 Laurson, L. et al. Evolution of the average avalanche shape with the universality class. Nat Commun 4, 2927 (2013).

33 Papanikolaou, S. et al. Universality beyond power laws and the average avalanche shape. Nat. Phys. 7, 316–320 (2011).

34 Sethna, J. P., Dahmen, K. a. & Myers, C. R. Crackling noise. Nature 410, 242–250 (2001).

35 Friedman, N. et al. Universal critical dynamics in high resolution neuronal avalanche data. Phys. Rev. Lett. 108, 208102 (2012).

36 Shaukat, A. & Thivierge, J.-P. Statistical evaluation of waveform collapse reveals scale-free properties of neuronal avalanches. Front. Comput. Neurosci. 10 (2016).

37 Dalla Porta, L. & Copelli, M. Modeling neuronal avalanches and long-range temporal correlations at the emergence of collective oscillations: Continuously varying exponents mimic M/EEG results. PLoS Comput. Biol. 15, e1006924 (2019).

38 Touboul, J. & Destexhe, A. Power-law statistics and universal scaling in the absence of criticality. Phys. Rev. E 95 (2017).

39 di Santo, S., Burioni, R., Vezzani, A. & Muñoz, M. A. Self-organized bistability associated with first-order phase transitions. Phys. Rev. Lett. 116, 240601 (2016).

40 Friedman, E. J. & Landsberg, A. S. Hierarchical networks, power laws, and neuronal avalanches. Chaos 23 (2013).

41 Roberts, J. A., Iyer, K. K., Finnigan, S., Vanhatalo, S. & Breakspear, M. Scale-free bursting in human cortex following hypoxia at birth. J. Neurosci. 34, 6557–6572 (2014).

42 Iyer, K. K. et al. Cortical burst dynamics predict clinical outcome early in extremely preterm infants. Brain 138, 2206–2218 (2015).

43 Karimipanah, Y., Ma, Z., Miller, J.-e. K., Yuste, R. & Wessel, R. Neocortical activity is stimulus- and scale-invariant. PLoS One 12, e0177396 (2017).

44 Buzsáki, G., Logothetis, N. & Singer, W. Scaling brain size, keeping timing: Evolutionary preservation of brain rhythms. Neuron 80, 751–764 (2013).

45 Lundqvist, M. et al. Gamma and beta bursts underlie working memory. Neuron 90, 152–164 (2016).

46 Engel, A. K. & Fries, P. Beta-band oscillations—signalling the status quo? Curr. Opin. Neurobiol. 20, 156–165 (2010).

47 Lisman, J. E. & Jensen, O. The theta-gamma neural code. Neuron 77, 1002–1016 (2013).

48 Nikolic, D., Fries, P. & Singer, W. Gamma oscillations: precise temporal coordination without a metronome. Trends Cogn. Sci. 17, 54–55 (2013).

49 Iemi, L. et al. Multiple mechanisms link prestimulus neural oscillations to sensory responses. eLife 8, e43620 (2019).

50 di Santo, S., Villegas, P., Burioni, R. & Muñoz, M. A. Landau–Ginzburg theory of cortex dynamics: Scale-free avalanches emerge at the edge of synchronization. Proc. Natl. Acad. Sci. U.S.A. 115, E1356–E1365 (2018).

51 Virkar, Y. & Clauset, A. Power-law distributions in binned empirical data. Ann. Appl. Stat. 8, 89–119 (2014).

52 Klaus, A., Yu, S. & Plenz, D. Statistical analyses support power law distributions found in neuronal avalanches. PLoS One 6, e19779 (2011).

53 Pritchard, W. S. The brain in fractal time: 1/f-like power spectrum scaling of the human electroencephalogram. Int. J. Neurosci. 66, 119–129 (1992).

54 Lundqvist, M., Herman, P., Warden, M. R., Brincat, S. L. & Miller, E. K. Gamma and beta bursts during working memory readout suggest roles in its volitional control. Nat. Commun. 9, 394 (2018).

55 Yu, S., Klaus, A., Yang, H. & Plenz, D. Scale-invariant neuronal avalanche dynamics and the cut-off in size distributions. PLoS One 9, e99761 (2014).

56 Priesemann, V., Valderrama, M., Wibral, M. & Le Van Quyen, M. Neuronal avalanches differ from wakefulness to deep sleep - Evidence from intracranial depth recordings in humans. PLoS Comput. Biol. 9, e1002985 (2013).

57 Priesemann, V., Munk, M. H. J. & Wibral, M. Subsampling effects in neuronal avalanche distributions recorded in vivo. BMC Neurosci. 10, 40 (2009).

58 Chen, Y. J., Papanikolaou, S., Sethna, J. P., Zapperi, S. & Durin, G. Avalanche spatial structure and multivariable scaling functions: Sizes, heights, widths, and views through windows. Phys. Rev. E 84, 061103 (2011).

59 Buzsáki, G. Rhythms of the Brain. (Oxford University Press, 2009).

60 Roux, F. & Uhlhaas, P. J. Working memory and neural oscillations: alpha–gamma versus theta–gamma codes for distinct WM information? Trends Cogn. Sci. 18, 16–25 (2014).

61 Fries, P., Nikolic, D. & Singer, W. The gamma cycle. Trends Neurosci. 30, 309–316 (2007).

62 Ermentrout, G. B. & Kleinfeld, D. Traveling electrical waves in cortex: insights from phase dynamics and speculation on a computational role. Neuron 29, 33–44 (2001).

63 Gireesh, E. D. & Plenz, D. Neuronal avalanches organize as nested theta- and beta/gamma-oscillations during development of cortical layer 2/3. Proc. Natl. Acad. Sci. U.S.A. 105, 7576–7581 (2008).

64 Lombardi, F., Herrmann, H. J., Plenz, D. & De Arcangelis, L. On the temporal organization of neuronal avalanches. Front. Syst. Neurosci. 8, 204 (2014).

65 Wang, S.-J. et al. Stochastic oscillation in self-organized critical states of small systems: Sensitive resting state in neural systems. Phys. Rev. Lett. 116, 018101 (2016).

66 Markram, H. et al. Reconstruction and Simulation of Neocortical Microcircuitry. Cell 163, 456–492 (2015).

67 Poil, S.-S., Hardstone, R., Mansvelder, H. D. & Linkenkaer-Hansen, K. Critical-state dynamics of avalanches and oscillations jointly emerge from balanced excitation/inhibition in neuronal networks. J. Neurosci. 32, 9817–9823 (2012).

68 Benayoun, M., Cowan, J. D., van Drongelen, W. & Wallace, E. Avalanches in a stochastic model of spiking neurons. PLoS Comput. Biol. 6, 21 (2010).

69 Priesemann, V. et al. Spike avalanches in vivo suggest a driven, slightly subcritical brain state. Front. Syst. Neurosci. 8, 108 (2014).

70 Fontenele, A. J. et al. Criticality between Cortical States. Phys. Rev. Lett. 122, 208101, (2019).

71 Ribeiro, T. L., Ribeiro, S., Belchior, H., Caixeta, F. & Copelli, M. Undersampled critical branching processes on small-world and random networks fail to reproduce the statistics of spike avalanches. PLoS One 9, doi:10.1371/journal.pone.0094992 (2014).

72 Chialvo, D. et al. How we move is universal: scaling in the average shape of human activity. Papers in Physics 7, 070017 (2015).

73 Proekt, A., Banavar, J. R., Maritan, A. & Pfaff, D. W. Scale invariance in the dynamics of spontaneous behavior. Proc. Natl. Acad. Sci. U.S.A. 109, 10564–10569 (2012).

74 Baldassarri, A., Colaiori, F. & Castellano, C. Average shape of a fluctuation: Universality in excursions of stochastic processes. Phys. Rev. Lett. 90, 060601 (2003).

75 Poil, S. S., Van Ooyen, A. & Linkenkaer-Hansen, K. Avalanche dynamics of human brain oscillations: Relation to critical branching processes and temporal correlations. Hum. Brain Mapp. 29, 770–777 (2008).

76 Wang, S.-J., Hilgetag, C. & Zhou, C. Sustained activity in hierarchical modular neural networks: self-organized criticality and oscillations. Front. Comput. Neurosci. 5 (2011).

77 Petkoski, S., Palva, J. M. & Jirsa, V. K. Phase-lags in large scale brain synchronization: Methodological considerations and in-silico analysis. PLoS Comput. Biol. 14, e1006160 (2018).

78 Deco, G. & Jirsa, V. K. Ongoing cortical activity at rest: criticality, multistability, and ghost attractors. J. Neurosci. 32, 3366–3375 (2012).

79 Ramos, O., Altshuler, E. & Måløy, K. Avalanche prediction in a self-organized pile of beads. Phys. Rev. Lett. 102, 078701 (2009).

80 Palva, S. & Palva, J. M. Roles of brain criticality and multiscale oscillations in temporal predictions for sensorimotor processing. Trends Neurosci. 41, 729–743 (2018).

81 Hidalgo, J. et al. Information-based fitness and the emergence of criticality in living systems. Proc. Natl. Acad. Sci. U.S.A. 111, 10095–10100 (2014).

82 Uhlhaas, P. J. & Singer, W. Neuronal dynamics and neuropsychiatric disorders: toward a translational paradigm for dysfunctional large-scale networks. Neuron 75, 963–980 (2012).

83 Seshadri, S., Klaus, A., Winkowski, D. E., Kanold, P. O. & Plenz, D. Altered avalanche dynamics in a developmental NMDAR hypofunction model of cognitive impairment. Translational Psychiatry 8, 3 (2018).

84 Yang, H., Shew, W. L., Roy, R. & Plenz, D. Maximal variability of phase synchrony in cortical networks with neuronal avalanches. J. Neurosci. 32, 1061–1072 (2012).

