## Supplemental Info for "The scale-invariant, temporal profile of neuronal avalanches in relation to cortical γ–oscillations"

Miller et al.

### Supplementary Tables and Figures

**Supplementary Table 1**: Summary of NHP LFP recordings analyzed in the current study. *Recording Time*: total duration of recordings concatenated over multiple recording sessions separated by up to several days. *Recording Sessions*: number of resting-state recording sessions. *Recording Span*: number of weeks over which the session data were collected. *Working Electrodes*: number of electrodes showing adequate SNR (<~7 s.d.) out of 96 electrodes on the array. *Artifact removal*: percentage of recording time removed due to artifacts caused by i.e. vocalization, sudden movements, chewing, etc. *s.d.*: standard deviation of post-cleaning channel activity, averaged over all working electrodes. *Mx*: NHP x. *PM*: premotor cortex. *PF*: prefrontal cortex. NHP K had two arrays in PF, whereas NHP V & N had 1 array in PM and PF each.

| | Recording Time (*min*) | Recording Sessions (*n*) | Recording Span (*wks*) | Functioning Electrodes (*n*) | Artifact Removal (%) | s.d. ($\mu V$) |
|---|---|---|---|---|---|---|
| V-PF | 167 | 6 | 2 | 91 | 2.5 | 25 |
| V-PM | 208 | 7 | 2 | 91 | 5.3 | 35 |
| N-PF | 135 | 7 | 8 | 89 | 9.9 | 35 |
| N-PM | 122 | 4 | 6 | 71 | 6.0 | 35 |
| K-PF1 | 562 | 19 | 11 | 81 | 7.0 | 140 |
| K-PF2 | 118 | 4 | 1 | 91 | 9.2 | 115 |
| mean ± s.d. | 219 ± 171 | 6 ± 6 | 8 ± 4 | 85 ± 8 | 7 ± 3 | 64 ± 50 |

**Supplementary Table 2**: Average nLFP event statistics on the array is similar across NHP and cortical area examined (thresholded at -2 s.d.). *Average Inter-Event Interval* ⟨IEI⟩: average duration of silence between successive suprathreshold (-2 s.d.) nLFPs on the array binned at 2 kHz sampling frequency. *Event Rate*: frequency at which suprathreshold nLFPs were detected. For legend see also Supplementary Table 1.

| | ⟨IEI⟩ (*ms*) | Event Rate (*Hz*) | $\langle IEI \rangle_{1\text{-}20\ Hz}$ (*ms*) |
|---|---|---|---|
| V-PF | 2.42 | 4.61 | 9.97 |
| V-PM | 3.60 | 3.02 | 16.00 |
| N-PF | 4.00 | 2.59 | 16.42 |
| N-PM | 3.90 | 3.55 | 16.28 |
| K-PF1 | 3.52 | 2.96 | 13.32 |
| K-PF2 | 2.88 | 3.75 | 11.61 |
| mean ± s.d. | 3.39 ± 0.62 | 3.37 ± 0.34 | 13.9 ± 2.7 |

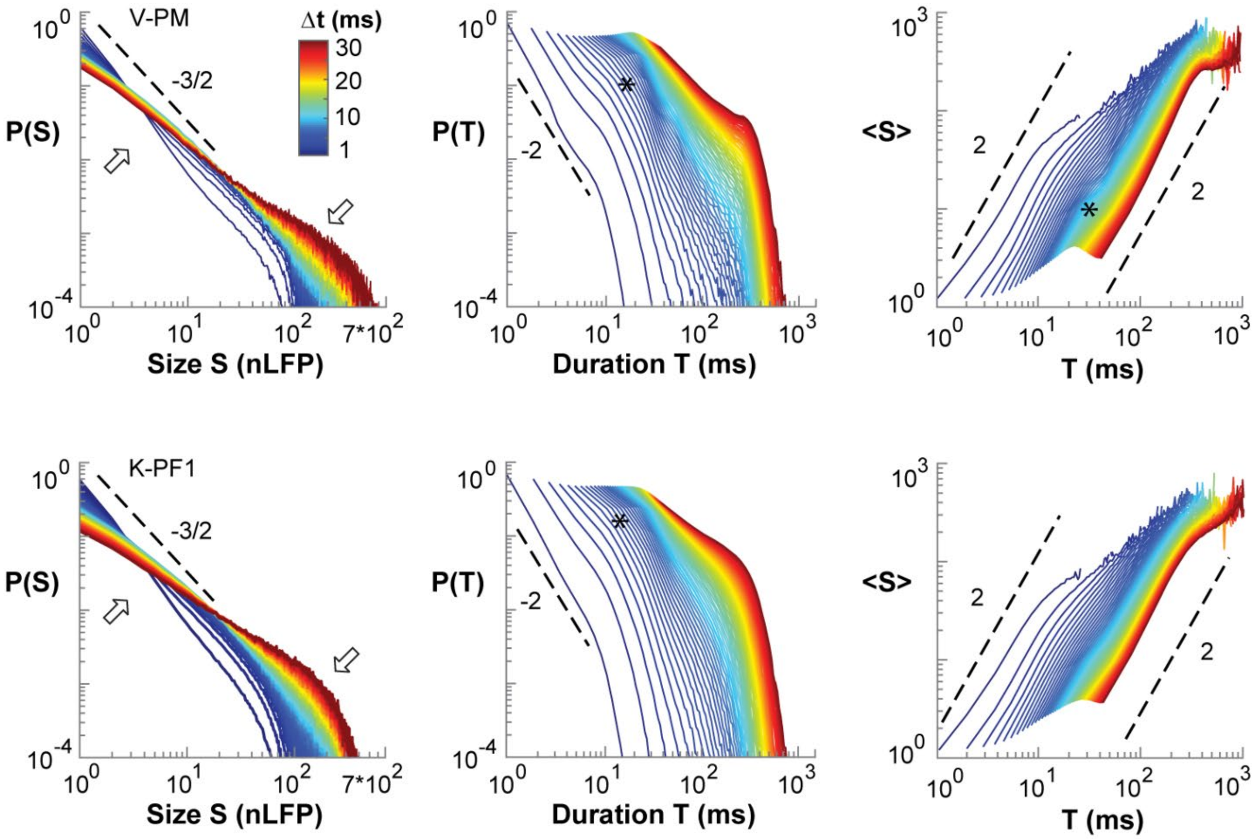


**Supplementary Fig. 1:** γ–oscillations and neuronal avalanches coexist in cortical resting activity of nonhuman primates. Power law in nLFP cluster sizes identifies avalanche dynamics for monkeys V-PM (*top*) and K-PF1 (*bottom*). Note cut-off at ~100 electrodes (*arrow*). *Dashed lines*: visual guides for power law slopes.

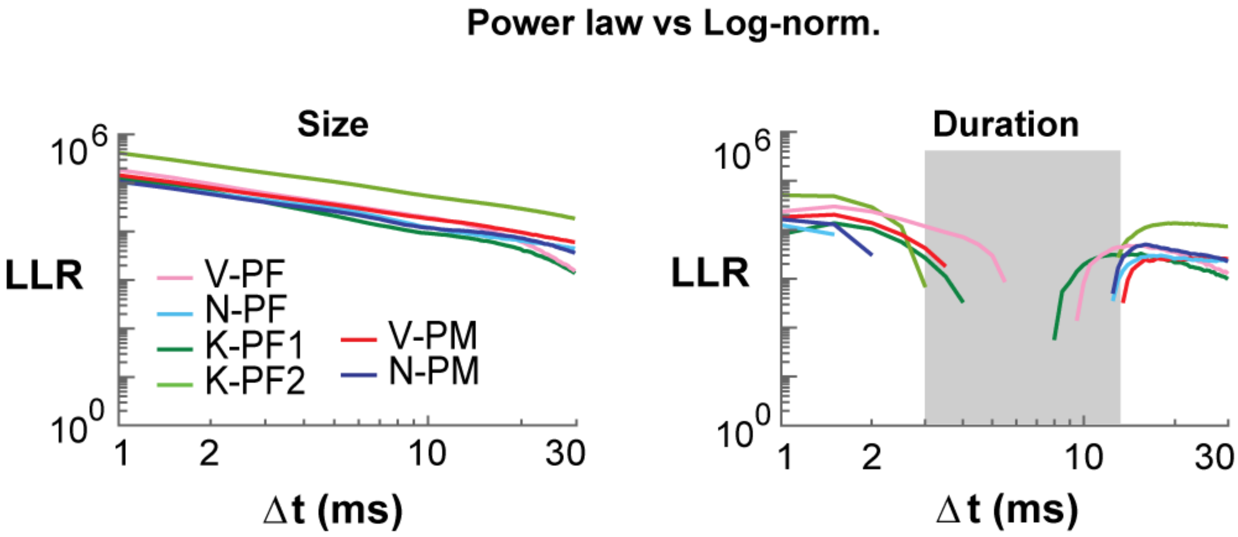


**Supplementary Fig. 2:** Log-likelihood test of neuronal avalanche size and duration distributions for 1–100 Hz LFP. Log-likelihood ratio comparing power law vs. log-normal fit to the avalanche size (*top*) and avalanche duration (*bottom*) distributions for decreasing temporal resolution $\Delta t$ in all arrays (for color code see Supplementary Fig. 1). Note LLR < 0 for lifetime distributions identifies a range $\Delta t$ = 3–15 ms, for which durations deviate from a power law ($P$ > 0.1) with corresponding undefined power law slope $\beta$ (*grey area*).

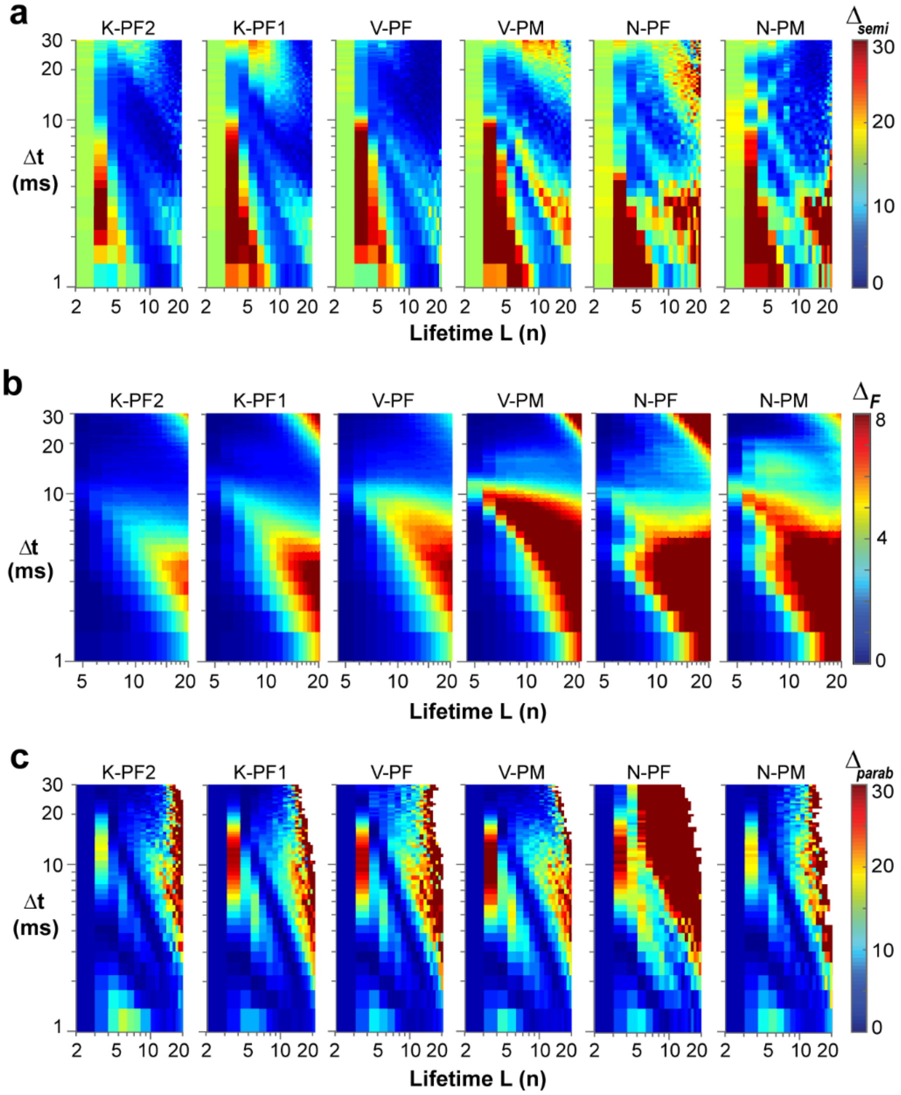


**Supplementary Fig. 3:** Supplemental density plots. **a**, Density plot of fit quality to a semicircle for all arrays and all profiles in the (L, Δt)-plane for LFP filtered from 1–100 Hz. Note consistent high error across the plane. **b**, Density plot of global (L = 3, …, 20) collapse error for arrays ranked from low (*left*) to high γ–oscillation power (*right*). Increase in γ–power leads to increasingly larger areas of deviation from good collapse.

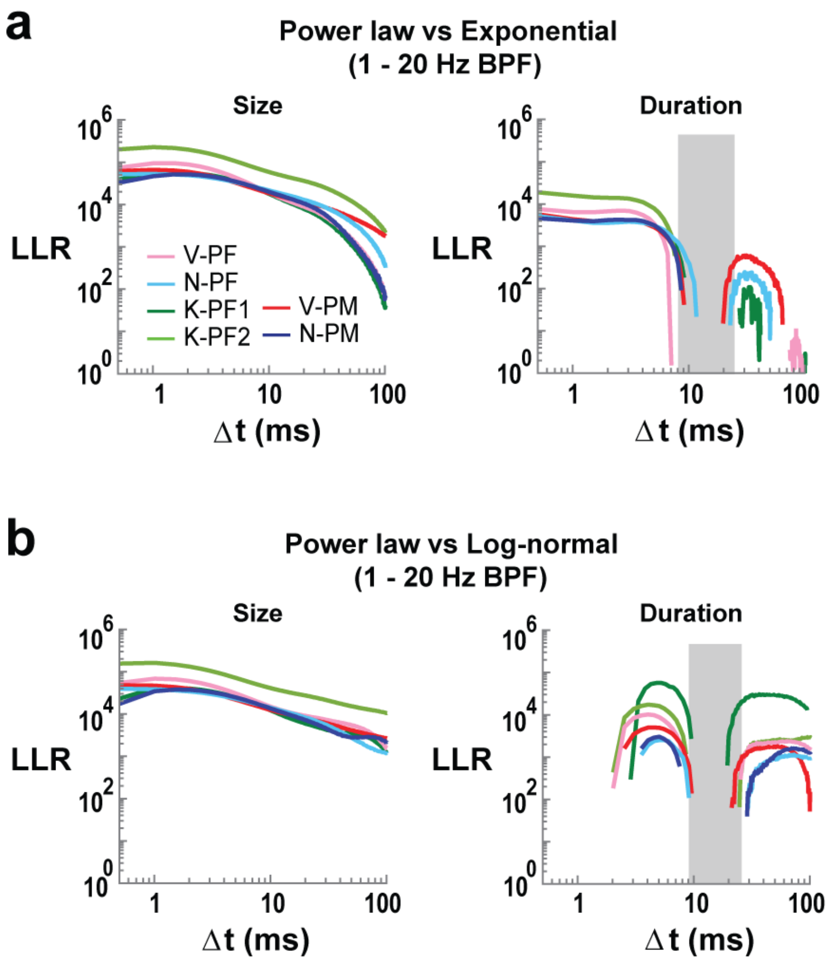


**Supplementary Fig. 4:** Log-likelihood test of neuronal avalanche size and duration distributions for 1–20 Hz LFP. **a**, Log-likelihood ratio (LLR) comparing power law vs. exponential model fits to the avalanche size (*top*) and avalanche duration (*bottom*) distributions for decreasing temporal resolution in all arrays. Note LLR of avalanche lifetime distributions reveals a range of $\Delta t = 8$–25 ms for which durations deviate from a power law (LLR < 0) and the slope $\beta$ is ill-defined (*grey area*). **b**, Corresponding comparison of power law vs. log-normal distribution.
